# Opportunistic evidence of the impact of bacterial infections on social integration in vampire bats

**DOI:** 10.1101/2023.04.17.537180

**Authors:** Imran Razik, Gerald G. Carter, Michael Abou-Elias, Sebastian Stockmaier

## Abstract

Social integration can affect an individual’s susceptibility to infectious disease. Conversely, infectious disease can reduce an individual’s social activity. Yet, it remains unclear to what extent short-term infections can inhibit social integration and the formation of new relationships. During a previous study on relationship formation, we captured 21 female common vampire bats (*Desmodus rotundus*) from different wild populations and housed them together in captivity. Upon introduction, we observed an unplanned outbreak of bacterial infections that caused cutaneous lesions. After treating infected bats with an antibiotic, five bats recovered, but four others suffered lasting injuries. Given that observations of how natural infections alter relationship formation are rare, we analyzed how allogrooming rates changed over time among familiar and new dyads consisting of the nine infected and 12 asymptomatic bats. We found that (1) infected bats demonstrated reduced activity and social behavior, (2) more severely infected bats gave and received less allogrooming than asymptomatic bats, (3) the effect of infection was larger for new dyads relative to familiar dyads, and (4) this effect decreased as infected bats recovered and new dyads became more familiar. These opportunistic observations were consistent with the hypothesis that short-term infections can impact the formation of new relationships.

---

Social integration, or the quantity and quality of an individual’s social relationships, can affect an individual’s susceptibility to infection (e.g., Pavez-Fox et al., 2021; Rushmore et al., 2017; Snyder-Mackler et al., 2016, 2020; Tung et al., 2012) and ability to tolerate or suppress the adverse effects of parasites (Almberg et al., 2015; Jog et al., 2022; Loehle, 1995; Stockmaier et al., 2023). Conversely, infectious disease can inhibit the social behaviors of infected individuals (Hart, 1988; Lopes et al., 2021; Stockmaier et al., 2021) by diminishing their motivation to interact with others (e.g., Fishkin & Winslow, 1997; Moreno et al., 2021; Stockmaier et al., 2018), reducing their movement (e.g., in mice (*Mus musculus domesticus*, Lopes et al., 2016) and mongoose (*Mungos mungo*, Fairbanks et al., 2014)), or causing others to avoid them (Behringer et al., 2006; Boillat et al., 2015; Hawley et al., 2020; Kiesecker et al., 1999; Lopes et al., 2022; Poirotte et al., 2017). These feedbacks between social integration and infectious disease are complex in part because the effects of infection on social behavior vary widely across pathogens, hosts, host sex, levels of infection severity, and types of social relationships (reviewed in Hawley et al., 2020). In particular, stable social bonds may be more important and should, therefore, be more resilient to immune challenges or changes in individual behaviors compared to weak social relationships (Poirotte & Charpentier, 2020; Seyfarth & Cheney, 2011; Stockmaier et al., 2020). For example, in vampire bats, sickness behavior reduces individual activity and social grooming among non-kin, but it does not decrease maternal grooming of offspring (Stockmaier et al., 2020). Similarly, mandrills avoid grooming parasite-infected non-kin but continue to groom infected close maternal kin (Poirotte & Charpentier, 2020). While these studies focus on the immediate effect of infections on social behaviors, it remains unclear to what extent infections impact affiliative social bond formation.

New social bonds form when individuals join a new social community, which is also when individuals are exposed to new pathogens. If exposure to new pathogens is more likely when encountering new social partners, and if infectious disease causes sickness behavior that limits individual activity and the motivation to interact, then acute infection during the earliest stages of relationship formation might negatively affect rates of social integration. In a group with both familiar and new social partners, we therefore predicted that the effect of infection on social integration should be stronger in new, unfamiliar dyads (pairs) compared to familiar dyads with well-established relationships. In new dyads of unfamiliar bats, infections should initially reduce social interaction rates for infected bats; as the infected individuals recover and sickness behaviors, such as lethargy and social withdrawal, resolve, interaction rates of infected and asymptomatic bats should then converge over time. In familiar dyads, we expected this interaction to be weaker because infections should have a smaller impact on how interaction rates change over time in more established relationships.

To address this hypothesis, we report opportunistic evidence that an acute bacterial infection reduced social integration in common vampire bats (*Desmodus rotundus*) in a captive setting. Social integration is observable in this species as strangers transition from no-cost clustering (huddling for warmth) to low-cost allogrooming (cleaning the fur or wings of other bats) and to high-cost food sharing (regurgitations of ingested blood to unfed individuals, Carter et al., 2020). Allogrooming is the best measure of relationship formation because it is a directed investment of time and energy (unlike clustering) that also occurs frequently enough to track over time through relationship formation (unlike food sharing). During a previous 4-month study, we tracked changes in allogrooming among 21 female vampire bats before and after a week of forced proximity in which small groups of three bats from three distant wild sites were housed together in captivity for one week (Razik et al., 2022). Shortly after the 21 bats from different sites were first introduced in captivity, we observed an unplanned outbreak of a bacterial infection, which we treated with antibiotics (Razik et al., 2022). In this paper, we re-analyzed published data on relationship formation by including new measures of infection severity to estimate the social effect of these infections. Specifically, we tracked changes in allogrooming over time among the nine infected female bats and the 12 asymptomatic female bats, comparing dyads that were previously familiar or unfamiliar with each other.

## Methods

### Ethics Statement

This work was reviewed and approved by the Smithsonian Tropical Research Institute Animal Care and Use Committee (STRI ACUC no. 2015-0501-2022) and the Panamanian Ministry of the Environment (no. SEX/A-67-2019).

### Study subjects

For an experiment testing the effects of forced proximity on relationship formation (Razik et al., 2022), we captured eight or nine common vampire bats from each of three sites in Panama that were 120-350 km apart (Figure 1). Upon capture, the 26 bats (23 adult females, two juvenile males, and one juvenile female) appeared healthy with no pathological lesions. We housed the bats together in an outdoor flight cage (2.1 × 1.7 × 2.3 m) in Gamboa, Panama, for approximately four months. Although we caught 23 females, two recaptured females entered the captive colony several weeks after the colony was created, were not included in all phases of our experiment, and were therefore not included in this analysis. We therefore analyzed the allogrooming relationships among 252 new and 126 familiar dyads of 21 adult females. We define ‘dyads’ as unique directed combinations of a groomer and receiver. We use the label ‘new’ for dyads introduced in captivity and ‘familiar’ for dyads captured from the same site (hollow tree or cave). Details of the study colony can be found in (Razik et al., 2022; Razik et al., 2021).

**Figure 1.**
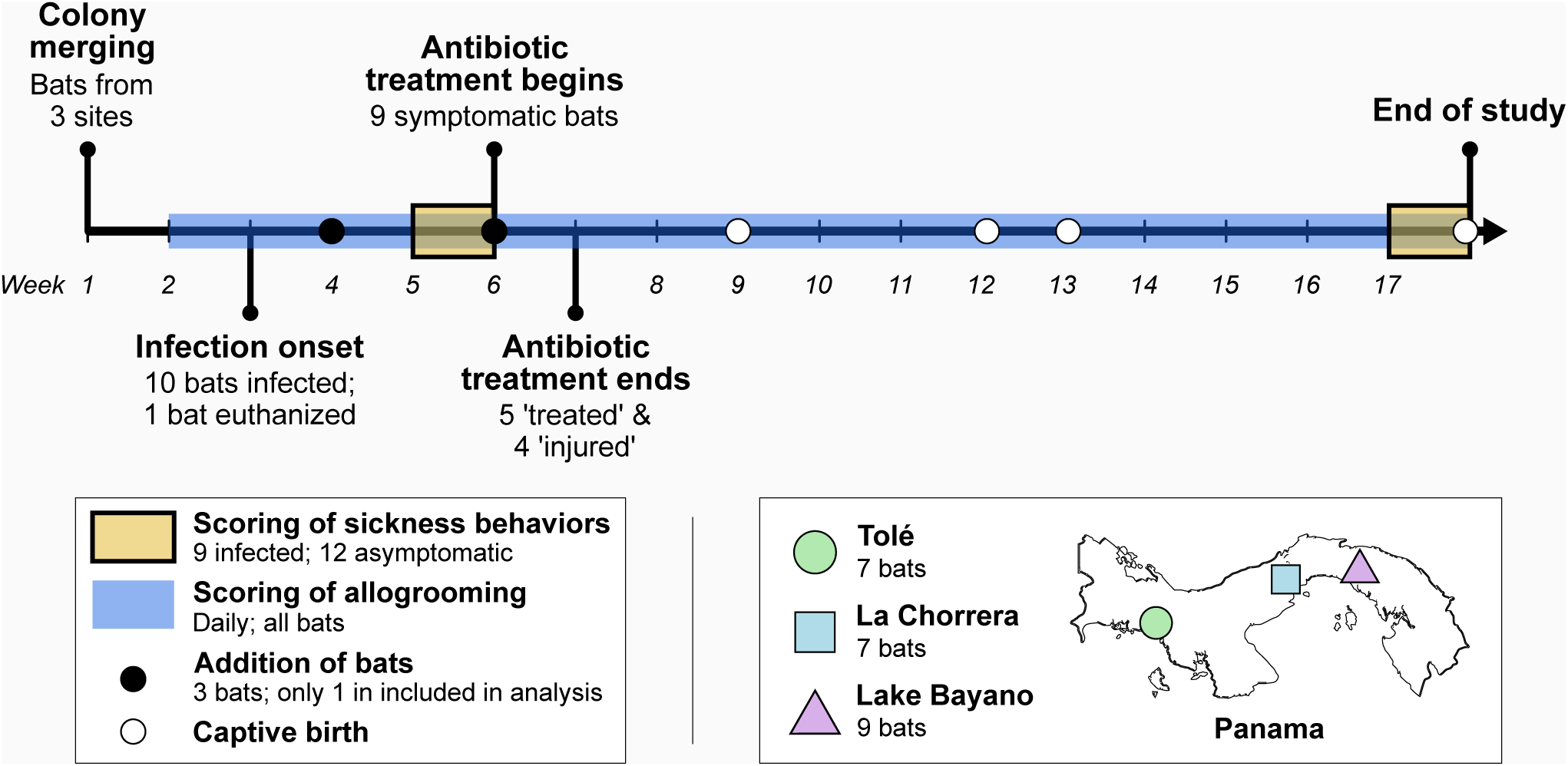
Detailed timeline of events.

### Behavior sampling

We used three infrared surveillance cameras to sample social interactions among bats individually marked with unique combinations of forearm bands (Razik et al., 2022). Over 114 days, we sampled 682 h of interactions, recording all visible allogrooming and mouth-licking interactions (≥ 5 seconds in duration) as well as any aggression. For each interaction, we noted the actor, receiver, and duration. We used allogrooming to assess social integration because allogrooming was 94% of the 17,592 observed social interactions, while mouth-licking and aggression were only 6% and 0.3% of interactions, respectively. For each directed edge, we summed the total seconds of allogrooming from the groomer to the receiver, and we summed the total sampled seconds that each dyad had the opportunity to interact. For further details on behavior sampling, see (Razik et al., 2022).

### Bacterial infection and treatment

Two weeks after the 26 wild-caught bats were first introduced to one another in the flight cage, 10 female bats began to show progressive symptoms of a cutaneous infection (blisters or open lesions with pus, as well as inflamed tissue). Symptoms progressed for approximately two weeks without signs of recovery. These infections were unexpected and, to our knowledge, have not previously been described in vampire bat colonies. The 10 infected bats varied in the severity of their symptoms. The most severe symptoms occurred in bats from a hollow tree near La Chorrera, Panama. The most severely infected bat developed large, pus-filled lesions on its back and had extreme inflammation around the wrists that led tissue to break apart and expose bone; we euthanized this bat before it received antibiotic treatment and did not include it in our analysis of social integration. Four other bats developed severe symptoms with lesions and inflammation on one or both wrists, leading to tissue scarring and the inability to sustain flight. One bat from the same roost experienced only minor inflammation and lesions on both wrists but healed without permanent tissue scarring and continued to fly. Three bats from a cave near Lake Bayano, Panama, were also infected. Two bats developed pus-filled abscesses: on the lower abdomen and upper thigh for one bat, on the lower back for the other bat, and the third bat developed only minor inflammation on 1-2 toes. Finally, one bat from a hollow tree near Tolé, Panama, had a large, round lesion on its knee, but this was evident at the start of the study and did not change over time. This bat displayed no other signs of infection. With most cases having been observed in bats from either La Chorrera or Lake Bayano, it seems the bats from Tolé were most resistant to the pathogen.

Swab and tissue samples from skin lesions were cultured to isolate bacterial pathogens, which indicated the presence of *Bacillus sp.* and *Staphylococcus aureus*. There are several *Bacillus* species (e.g., *Bacillus anthracis, Bacillus cereus,* and *Bacillus pumilus*) that may infect the skin or colonize wounds, but these infections are uncommon and often present with classic symptoms (e.g., black eschars and necrosis; Cromartie et al., 1947; Esmkhani & Shams, 2022; Tena et al., 2007; Veysseyre et al., 2015) that we did not observe in our colony. Alternatively, several species within the *S. aureus* complex, including *Staphylococcus schweitzeri* and *Staphylococcus argenteus,* among others (Becker et al., 2019; Chew et al., 2021; Tong et al., 2015), typically infect the skin and cause symptoms consistent with what we observed. *S. aureus* complex has also been isolated from the feces of Old-World fruit bats (Fountain et al., 2022; Held et al., 2017; Monecke et al., 2022; Olatimehin et al., 2018) and the common vampire bat (Chaverri, 2006; Moreno et al., 1975).

A Kirby-Bauer antibiogram indicated the susceptibility of the isolated bacteria to the antibiotic Enrofloxacin. Given the urgency and results of the antibiogram, we immediately used Enrofloxacin antibiotic to treat the infected female bats (described above) over the course of seven days via daily subcutaneous injections, but we did not perform additional testing to identify the bacterial species or strain. After the antibiotic treatment of the remaining nine infected bats (one was euthanized, see above), a subset of five bats recovered without injury (henceforth “treated”), but the other four bats suffered lasting physical impairments that rendered them unable to fly for the remainder of the study (henceforth “injured”). We refer to the remaining 12 bats in our analyses as “asymptomatic.” For a detailed timeline of events, see Figure 1, and for details on experimental bats, see Table S1.

### Testing for sickness behavior

To confirm that bacterial infections reduced individual activity and social behavior, i.e., ‘sickness behavior’, we compared the activity of nine infected, symptomatic bats versus 12 asymptomatic bats (all adult females) during a sample period of 1 week before the antibiotic treatment (pre-treatment week) and again during 1 week at the end of the study (post-treatment week) (Figure 1). An observer scored videos using focal sampling for the presence or absence of three behavioral states: moving, (all movement, e.g., changing location in the cage), allogrooming (including mouth-licking), and self-grooming, following definitions in (Stockmaier et al., 2018). The observer could not be entirely blind to the condition of bats in the videos because lesions were visible, but the analyzed behavioral categories were not subjective. For one hour (starting one hour after sunset), an observer noted at 30 sec intervals whether a focal female was engaged in any of the behaviors of interest. Therefore, we sampled each of the 21 bats for 14 days, resulting in 1680 samples per bat (120 per day).

To test if infection predicted the frequency of each behavioral state, we fit Bayesian multilevel binomial regressions in R (brms package, Bürkner, 2017) and obtained median posterior parameter estimates and 90% highest posterior density intervals (HPDI), i.e., the most likely range of parameter values given the model. We used a 90% interval to prevent readers from interpreting HPDIs as 95% confidence intervals or denoting “statistical significance.” For each behavioral state, model predictors were infection status (asymptomatic vs. infected), treatment period (pre- vs. post-treatment), the interaction between infection status and period, and bat identity as a random intercept. The response variable was the proportion of observations doing a behavior (allogrooming, self-grooming, and moving) out of the total number of observations of the bat on camera. To evaluate interactions between infection and treatment period, we estimated marginal means using the *emmeans* package (Lenth & Piaskowski, 2025). Across models, all parameters had above 1000 effective samples and an R-hat equal to 1.00 with four chains, indicating successful convergence. Since the effects of acute bacterial infection on vampire bat behavior have not been observed, we used conservative default priors across all effects. However, to put our effect sizes in context, we compared them with effect sizes from past experimental immune challenges, explained below.

### Comparison of results to previous experimentally induced sickness behavior

We present our results alongside reference effect sizes from two past experiments on the behavioral responses of female vampire bats to a simulated bacterial infection (Stockmaier et al., 2018, 2020). In both experiments, behaviors of vampire bats were observed after being injected with either lipopolysaccharide (LPS) or saline. Injections of LPS mimic bacterial infections and cause physiological (e.g., increased white blood cell counts) and behavioral (e.g., lethargy, reduced grooming) symptoms in vampire bats (see Stockmaier et al., 2018). In the first study (Stockmaier et al., 2018), vampire bats were observed in groups of 2-4 bats within small cages. LPS injection reduced the odds of allogrooming by a factor of 3.73 (binomial mixed effect model, Odds Ratio, 95% confidence interval (CI) = 2.25–6.50) and reduced the odds of self-grooming by a factor of 16 (Odds Ratio, 95% CI = 12–27). In the second study (Stockmaier et al., 2020), vampire bats were fasted to induce food sharing and observed in a larger group (36 bats) in a flight cage. Under these conditions, LPS injection reduced the odds of allogrooming by a factor of 1.89 (Odds Ratio, 95% CI = 1.84–1.93). This smaller effect on allogrooming could be due to differences in group size, cage size, the hunger state of the injected bats, or some combination of those factors. We present reference effects from both studies because the present study shares some features of both; the bats were unfasted like the first study, but held in a large group in the same flight cage as the second study.

### Effects of infection on symmetry of allogrooming given and received

For the bats that we studied, we expected that any sickness effects on allogrooming given would cause at least some effects on allogrooming received because allogrooming is highly reciprocal, and most allogrooming occurs bidirectionally (Carter & Wilkinson, 2013; Imran Razik et al., 2022; Wilkinson, 1986). To assess whether infection reduced or removed the symmetry of directed allogrooming, we compared the proportion of hourly allogrooming rates that were bidirectional for dyads with at least one infected bat versus dyads with two asymptomatic bats.

### Effects of infection and familiarity on dyadic allogrooming rates over time

We assessed how allogrooming changed over time among familiar and new dyads of bats that were either asymptomatic, infected, or injured. To do this, we fit Bayesian social relation models in R (STRAND package, Ross et al., 2024). We used an aggregated binomial outcome to model the proportion of total seconds of observation time with or without allogrooming for each directed groomer-receiver dyad. Four dyadic covariates included whether the dyad had been forced into proximity (co-housed in a small cage) during a previous experiment (Razik et al., 2022, true/false), as well as the main effects and interaction between familiarity (true/false) and infection severity (a sum of both individuals’ infection severity scores: 0 = asymptomatic, 1 = infected, 2 = injured). This measure of infection severity assumes that these states have linearly increasing and additive effects, but this simplifying assumption is consistent with comparisons of allogrooming for infected bats versus all others (Fig. 2) and for injured bats versus all others (Fig. S1). To explore how these effects change through time, we binned the data by week (15 social networks across 16 weeks, excluding one week when bats were not sampled altogether), fit the same model described above to each time bin, and extracted posterior parameter estimates with 90% HPDIs. Model convergence diagnostics can be found in the online data repository (47).

**Figure 2.**
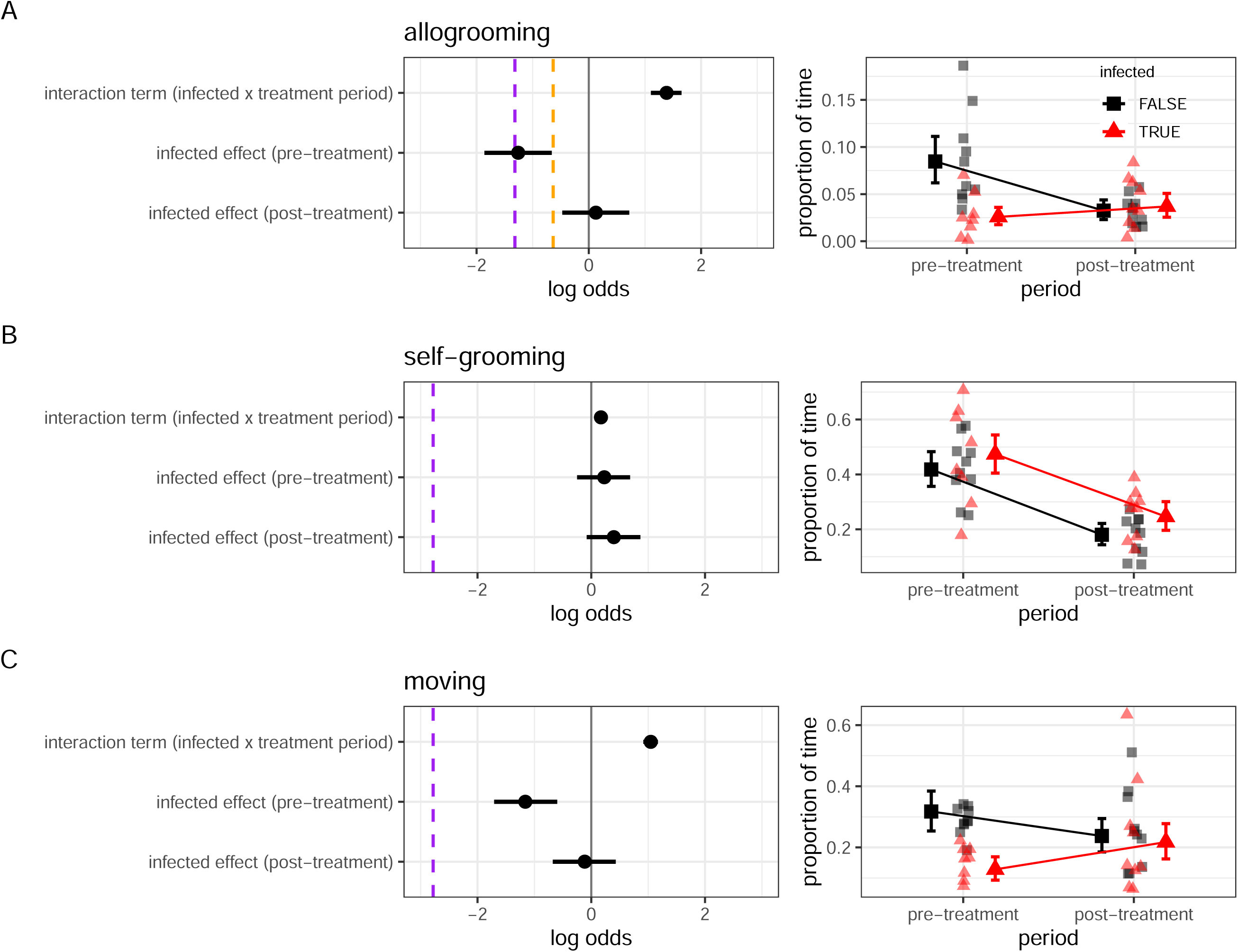
Infections reduced allogrooming and moving, but not self-grooming in vampire bats. *Note.* Figure panels show the effects of infections on allogrooming (A), self-grooming (B), and moving (C). Left-hand panels show median posterior parameter estimates (points) with 90% highest posterior density intervals (HPDI). Vertical dashed lines show previous effect sizes estimated from two LPS-induced immune challenge experiments (purple, (Stockmaier et al., 2018); orange, (Stockmaier et al., 2020)). Right-hand panels show proportions of time for each behavior (jittered data) for infected bats (red triangles) and asymptomatic bats (black squares) during the outbreak (pre-treatment) and after bats recovered (post-treatment). Points with error bars are predicted mean proportions with 90% credible intervals, marginalizing over random effects.

We predicted a three-way interaction between familiarity, time, and infection severity. Specifically, we expected that (1) infections would have a greater negative impact on allogrooming rates in new dyads relative to familiar dyads, and (2) the effect of infection on the slope of dyadic allogrooming rates over time would be greater for new dyads than for familiar dyads. Over time, both infection and familiarity effects should converge towards zero because by the end of the study, new dyads are no longer completely new, and infected bats are no longer infected.

## Results

### Confirming sickness behavior

Consistent with sickness behavior and recovery, infected bats spent less time moving and allogrooming than asymptomatic bats before antibiotic treatment. These differences faded to zero over time (Figure 2A, Figure 1C). Before antibiotic treatment, infected bats spent 2.6% of their time allogrooming, 90% credibility intervals = [1.8, 3.6%], whereas asymptomatic bats spent 8.4% [6.1, 11.1%] of their time allogrooming. Infected bats spent 12.8% [9.3, 16.9%] of their time moving, compared to 32.0% [25.9, 38.7%] in asymptomatic bats. After recovery, the previously infected bats no longer showed differences from the asymptomatic bats in allogrooming or movement (Figure 2A, Figure 2C). In contrast with effects on allogrooming, infections did not clearly reduce self-grooming (Figure 2B).

### Effects of infection on symmetry of allogrooming given and received

We found no evidence that infection reduced or removed allogrooming symmetry. Dyadic allogrooming was bidirectional within the hour in 84% of cases with at least one infected bat and in 81% of cases with two asymptomatic bats. The observation that allogrooming with an infected bat was not more likely to be unidirectional than allogrooming among asymptomatic bats confirms our assumption that any effect of infection on allogrooming given would also impact allogrooming received.

### Effects of infection and familiarity on dyadic allogrooming rates over time

The interaction between familiarity and infection severity changed over time (Figure 3). Prior to antibiotic treatment (weeks 1 – 6), estimates of the interaction between familiarity and infection severity were largely negative showing that the effect of infection was greater for new dyads than for familiar dyads (Figure 3). As expected, this difference then faded during the post-treatment period (weeks 8 – 16) as introduced bats became more familiar and as the infected bats recovered (Figure 3). In familiar dyads, infection did not clearly impact dyadic allogrooming rates over time, whereas in new dyads, there was a clear negative impact of infection severity on allogrooming rates that diminished with time (Figure 4).

**Figure 3.**
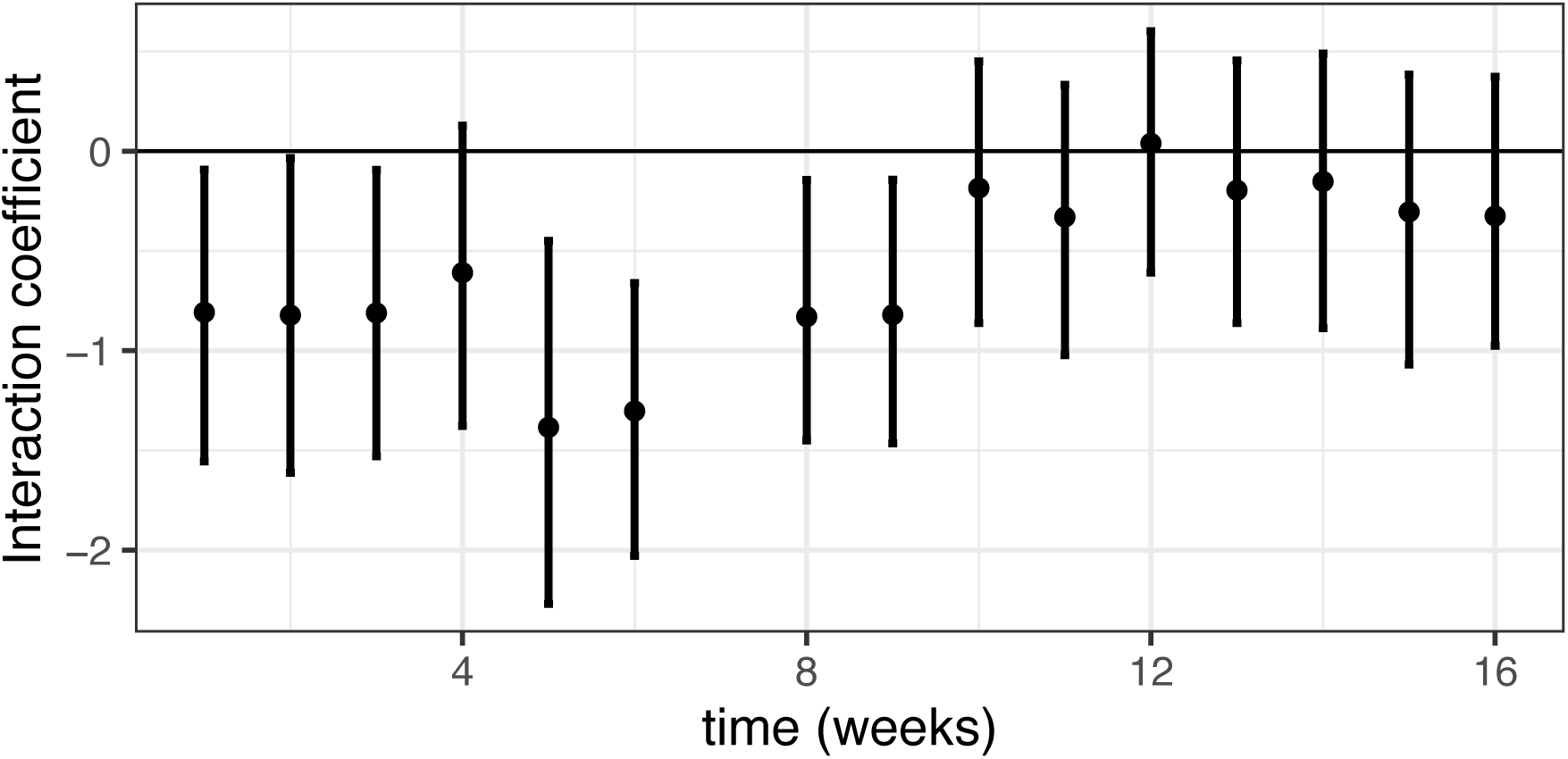
The effect of infection on allogrooming initially differs between new and familiar dyads, but this difference faded with time as bats recovered and became familiar. *Note.* Y-axis shows the interaction between familiarity and infection severity as predictors of allogrooming rate for each week (x-axis). Points with error bars show median posterior parameter estimates with 90% highest posterior density intervals.

**Figure 4.**
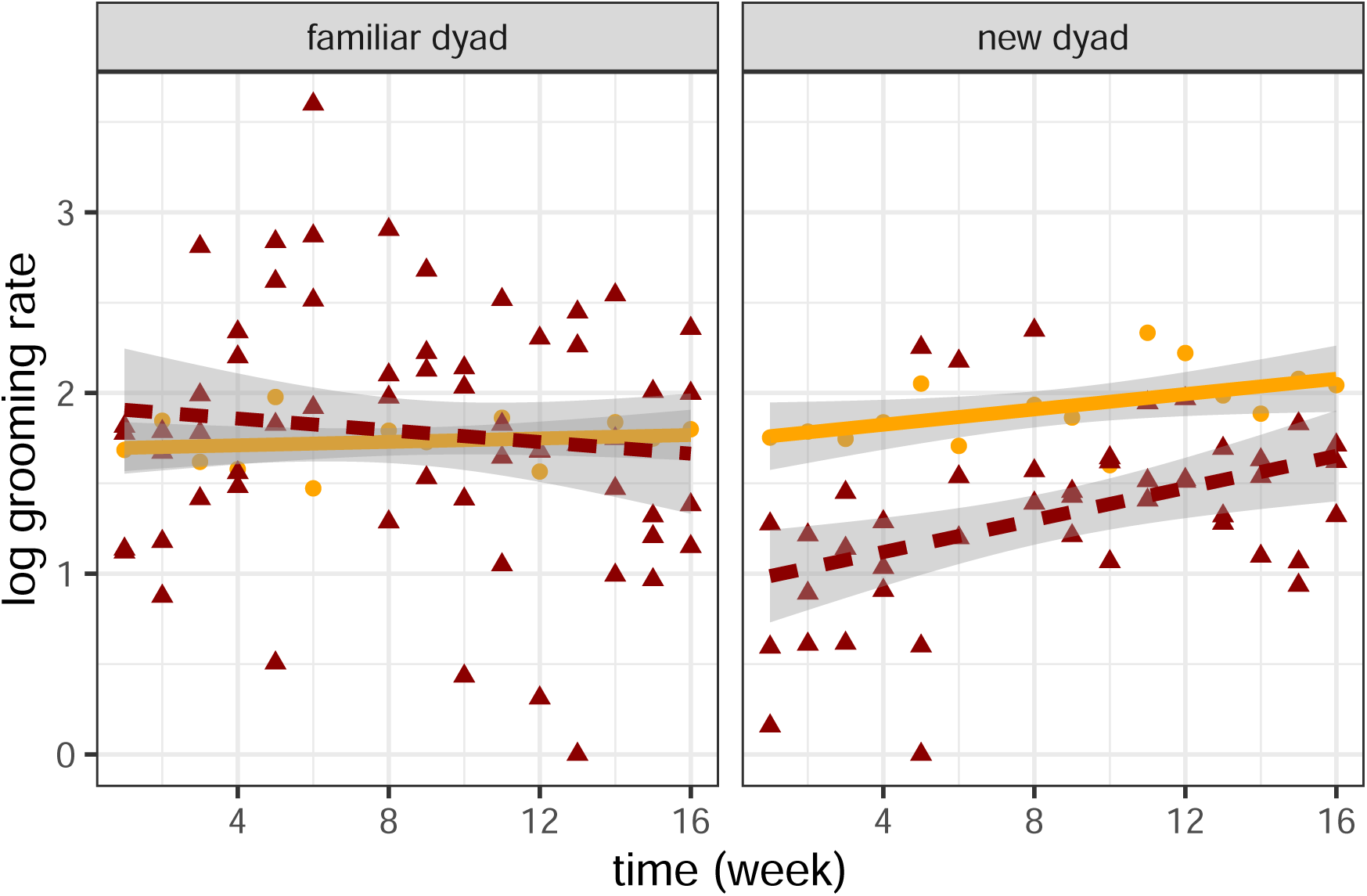
Evidence that infections disproportionately reduced allogrooming in new dyads (developing social relationships) relative to familiar dyads (existing social relationships). *Note.* Y-axis shows the log-transformed (log(x + 1)) mean daily allogrooming rates among all familiar dyads (left) and new dyads (right) across weeks. Yellow points and solid slopes indicate dyads with an infection severity of 0-1 (e.g., two asymptomatic bats, or one infected bat), whereas dark red triangles and dashed slopes indicate dyads with an infection severity of 2-4 (e.g., two infected bats, two injured bats). Shaded regions indicate 95% CI of the slope.

## Discussion

We documented opportunistic evidence that acute symptoms of bacterial infection can limit the development of allogrooming relationships in vampire bats. Relative to asymptomatic bats, the initially infected bats spent less time moving and allogrooming. This sickness behavior, which lasted approximately 2-3 weeks, led to changes in social integration, whereby infections had a greater impact on the formation of new allogrooming relationships than on allogrooming in existing relationships. Allogrooming rates were therefore influenced by a three-way interaction between familiarity, infection severity, and time. Infections had a more negative impact on allogrooming rates in new dyads relative to familiar dyads, such that the slope of allogrooming over time was greatest for new dyads with infected and/or injured bats. Familiar dyads and new dyads without infected bats were less impacted by infections. As predicted, the effects of infection and familiarity effects, as well as their interaction, converged towards zero as new dyads became familiar and infected individuals recovered over three months. This convergence suggests that sickness effects were not entirely driven by injuries that reduced movement.

Because we did not randomly and experimentally infect the bats, a possible alternative explanation for our findings is the reverse causal path to what we suggested above: perhaps bats with worse social integration (or worse social competence) had weaker immune systems and, therefore, suffered worse infections. However, this hypothesis cannot easily explain why the gap in social integration faded over time after antibiotic treatment. If the infected bats were already less social or less healthy for reasons besides the infection, then we would expect their social integration to be worse throughout the study, not just during the outbreak. Also, if social factors (like dominance or social competence) caused infection severity, then infection rates should be similar within each wild population. Instead, infection rate and severity were associated with wild population, suggesting that infections were a cause rather than a consequence of how bats formed new relationships.

The sickness behavior we observed was similar in some respects to the responses of vampire bats to simulated infections from injections of lipopolysaccharide, LPS (Stockmaier et al., 2018, 2020). Using immune challenges like LPS allows for controlled experiments on the connection between the inflammatory and the behavioral response to infection (Alciatore et al., 2021; Fishkin & Winslow, 1997; Lopes et al., 2012, 2016; Love et al., 2023; Moreno et al., 2021; Stockmaier et al., 2018, 2020; Willette et al., 2007), but naturally infecting pathogens can have different, pathogen-specific effects. For instance, LPS-injected vampire bats significantly reduced their self-grooming. Still, the naturally infected bats in our study did not, possibly because the bacterial pathogens caused skin lesions that encouraged self-grooming even when overall activity was reduced. Another pathogen that infects the skin of bats, the fungus *Pseudogymnoascus destructans* (the cause of ‘white-nose syndrome’) also causes increased self-grooming (Brownlee-Bouboulis & Reeder, 2013).

These observations are opportunistic and sparse (only 9 infected bats), but we believe our results are valuable for two main reasons. First, effect size estimates for sickness behaviors of vertebrates in response to naturally infecting pathogens are uncommon compared to experiments that use immune challenges (but see Bouwman & Hawley, 2010; Brownlee-Bouboulis & Reeder, 2013; Cárdenas-Canales et al., 2022; Fairbanks et al., 2015; Kollias et al., 2004; Langager et al., 2023). Second, while several studies have quantified the effect of immune challenges (infection or immunogenic antigens) on host social behavior (e.g., Fishkin & Winslow, 1997; Lopes et al., 2016; Moreno et al., 2021; Stockmaier et al., 2018; Willette et al., 2007), the effect of short-term infections on the formation and development of long-term social bonds remains largely unexplored. Our results can inform future experiments on the effect of infections on social integration and relationship formation.

Our findings also raise several questions. When might animals suppress their infection-induced behavioral symptoms to continue social integration (Aubert et al., 1997; Langager et al., 2023; Lopes, 2014; Lopes et al., 2012; Stockmaier et al., 2020; Weil et al., 2006; Willette et al., 2007)? At what stages of relationship formation do sickness behaviors have the most substantial effect on social interactions? What is the relationship between the duration or intensity of sickness behavior and the effects on social integration? To address these questions experimentally, researchers could use immune challenges at different doses (Dantzer et al., 1998; Grigoleit et al., 2011; Willette et al., 2007), and/or administer them at varying stages of relationship formation. Such standardized immune challenges may be better suited for isolating the effects of sickness on social integration because they minimize confounding variation in pathogen exposure or host vulnerability. In our case, differences in infection severity among source populations suggested that susceptibility might have depended on factors such as immune status or microbiome composition. For instance, site- or region-specific variation in the skin microbiota (as shown in other bat species, see Avena et. al., 2016) could have influenced colonization by opportunistic, pathogenic bacteria and the severity of resulting lesions, which, in turn, could have affected the duration of sickness behavior and opportunities for social integration.

In conclusion, we found opportunistic observational evidence for reduced social integration in infected vampire bats after an outbreak of a naturally infecting pathogen. Just as sickness behavior and other cues of infection can reduce social activity (reviewed in Hart, 1988; Loehle, 1995; Lopes et al., 2021; Stockmaier et al., 2021) and mate attraction (reviewed in Beltran-Bech & Richard, 2014), there may be similar effects on the formation of same-sex social bonds. If there are critical periods of relationship formation, then infections during this time might have lingering social consequences.

## Acknowledgments

The authors thank Rachel Page for lab space and resources, the staff at the Smithsonian Tropical Research Institute, and the following individuals who assisted with vampire bat care and/or data collection: D. Aparicio, B. Brown, G. Cohen, L. Dück, D. Girbino, E. Kline, C. Marroquin, and S. Ripperger. We thank two anonymous reviewers for their helpful comments.

## Author contributions

Conceptualization: IR, GC, SS; Data Curation: IR, GC; Formal Analysis: IR, GC, SS; Funding Acquisition: IR, GC; Investigation: IR, MA, GC; Methodology: IR, GC; Project Administration: IR, GC; Resources: IR, GC; Software: IR, GC; Supervision: GC, SS; Validation: IR, GC, SS; Visualization: IR, GC; Writing – Original Draft Preparation: IR, GC, SS; Writing – Review & Editing: IR, MA, GC, SS

## Data accessibility

Data and R code for all analyses are available on Github (Razik et al., 2023).

