## Supplementary material for "Opportunistic evidence of the impact of bacterial infections on social integration in vampire bats": Table S1

**Supplementary Materials**

**Table S1:** Summary of bats included in bacterial infection study. Severity rank reflects increasing severity of infection (higher values indicate more severe and persistent impairment).

| Bat ID | Bat Name | Capture Location | Capture Date | Infected | Severity Rank | Clinical Signs During Infection |
| --- | --- | --- | --- | --- | --- | --- |
| add | Lilith | Lake Bayano | 5 May 2019 | No | NA | None observed |
| addld | Queen | Lake Bayano | 5 May 2019 | No | NA | None observed |
| addldd | Irma | Lake Bayano | 5 May 2019 | No | NA | None observed |
| adld | Kelly | Lake Bayano | 5 May 2019 | Yes | 1 | Minor swelling of 1–2 toes; resolved rapidly following antibiotic treatment |
| adldd | Tiffany | Lake Bayano | 5 May 2019 | Yes | 4 | Single pus-filled lesion on ventral abdomen/upper thigh; healed well; limited mobility due to prior wing injury |
| aldd | Shania | Lake Bayano | 5 May 2019 | Yes | 3 | Single pus-filled lesion on dorsal lower back; healed without complications |
| ax | Jupiter | Lake Bayano | 5 May 2019 | No | NA | None observed |
| bdlx | Cindy | Tolé | 8 Jun 2019 | No | NA | None observed |
| bes | Esther | Tolé | 8 Jun 2019 | No | NA | None observed |
| bey | Betty | Tolé | 8 Jun 2019 | No | NA | None observed |
| blx | Gayle | Tolé | 8 Jun 2019 | Yes | 2 | Swollen knee present at study onset; minimal response to antibiotics |
| bs | Hera | Tolé | 8 Jun 2019 | No | NA | None observed |
| bxdlx | Fran | Tolé | 8 Jun 2019 | No | NA | None observed |
| bxx | Mabel | Tolé | 8 Jun 2019 | No | NA | None observed |
| cdw | Vixen | La Chorrera | 24 May 2019 | Yes | 5 | Bilateral wrist swelling; healed without permanent scarring; flight maintained |
| cfd | Zelda | La Chorrera | 24 May 2019 | Yes | 7 | Severe left wrist lesions; permanent scarring prevented sustained flight |
| cldw | Winona | La Chorrera | 24 May 2019 | Yes | 8 | Severe right wrist lesions; permanent scarring prevented sustained flight |
| clw | Rina | La Chorrera | 24 May 2019 | No | NA | None observed |
| clwd | Alicia | La Chorrera | 13 June 2019 | No | NA | None observed |
| cnone | Piper | La Chorrera | 24 May 2019 | Yes | 9 | Severe bilateral wrist lesions; permanent impairment of finger control and flight |
| cww | Opal | La Chorrera | 24 May 2019 | Yes | 6 | Severe left wrist lesions; permanent scarring prevented sustained flight |
